# A demand-based framework explains prioritization strategies upon transient limitations of different amino acids

**DOI:** 10.1101/2023.07.31.551408

**Authors:** Ritu Gupta, Swagata Adhikary, Nidhi Dalpatraj, Sunil Laxman

## Abstract

Cells require disparate amounts of distinct amino acids, which themselves have discrete biosynthetic costs. However, it remains unclear if and how cells respond differently to their scarcity. To explore this, we re-organized amino acids into distinct groups based on their metabolic origins. Subsequently, using yeast we assessed responses to transient disruptions in amino acid supply, and uncover diverse restoration responses for distinct amino acids. Cells hierarchically prioritize restoring glutamate-, sulfur-, pentose-phosphate- and pyruvate-derived amino acids. Particularly, the strongest response is to the glutamate-derived amino acid arginine. We find that the extent and priority of the restoration response is determined by the composite demand for an amino acid, coupled with low individual biosynthetic costs of that amino acid. We propose that cells employ a conserved strategy guided by the law of demand, to prioritize amino acids restoration upon transient limitation.

## Introduction

As environments change, cells sense the presence or absence of nutrients, and accordingly rewire metabolic processes towards restoring homeostasis. Cell metabolism can be viewed as an economy with the ability to manage metabolite supply in proportion to their requirements towards different allocations^1–3^. Such a ‘metabolic factory’ should not merely synthesize different metabolites to satisfy its activities, but ensure that each of these is supplied in the right quantities at the right time^4^. Cells match this required supply to demand through coordinated responses. Indeed, several studies have uncovered global metabolic programs that regulate growth in the context of different nutrient environments^5–9^. However, how cells might prioritize the restoration of distinct metabolic resources upon transient disruptions in their supply remains unresolved.

Amino acids form the core of a cellular economy, by being essential for protein synthesis as well as metabolism, and the scale of the latter is underappreciated. Given this, multiple mechanisms exist to sense amino acid sufficiency and restore amino acid balance. For example, in eukaryotic cells, the activity of the target of rapamycin complex 1 (TORC1) indicates the extent of amino acid demand for growth^10,11^. Contrastingly, the Gcn4/Atf4 transcription factor functions during starvation and growth, to respond when amino acid supply does not match demand, by driving amino acid biosynthesis to match demand^12–14^, thereby functioning as a sensor enabling metabolic control strategies^3^. A cell incapable of considering both the supply parameters as well as the demand for that molecule would struggle to survive^1, 2^. Surprisingly, in the context of amino acids, when the supply is disrupted (causing transient supply-demand mismatches), our knowledge of any prioritization strategies to restore distinct amino acids remains limited. Studies with complete amino acid starvations treat all amino acids fairly uniformly^6, 9,15^. However, amino acids each have distinct metabolic origins and synthesis routes^16, 17^, varying uses, and dissimilar intracellular concentrations^18^. It is currently unknown if cells treat temporary disruptions in the supply of any amino acid uniformly, or if they differently prioritize restoring the supply for distinct amino acids.

In this study, using prototrophic yeast cells in a defined glucose and nitrogen replete environment, and disrupting exogenous amino acid supply, we asked how cells prioritize restorations for distinct amino acids. We organized amino acids into groups based on their metabolic origins, transiently disrupted the supply of each group, and used reporters to assess amino acid supply-demand mismatches. Through this, we uncover hierarchically prioritized restoration responses when the supply of distinct amino acid group are transiently disrupted, with the strongest response towards the glutamate-derived amino acid arginine. The extent of restoration responses were consistent with a demand-driven economy, where the highest response observed correlated with a combination of low biosynthetic costs and high collective demand for that amino acid. These findings can be leveraged for the metabolic engineering of cells, and predict nutrient sensing responses.

## Results

### Cells exhibit hierarchical responses to supply-disruptions of amino acid groups, notably the glutamate-derived group

The supply of amino acids comes from extrinsic sources, as well as via *de novo* synthesis from carbon and nitrogen precursors. The demands for amino acids comes from protein synthesis, as well as their uses in metabolism^19, 20^, as illustrated in Fig1A. When there are disruptions in amino acid supply (e.g. by restricting extrinsic sources), we wondered how cells prioritized the restoration of distinct amino acids, by assessing changes in either demand or supply (Fig 1A). To better reflect supply criteria, we first organized amino acids into groups based on their metabolic origins. Conventional amino acid groupings are based on physicochemical properties (Fig S1A), without considering their metabolic origins. However, the ability of a cell to supply different precursors for a given amino acid is not uniform, depending on available precursors and the metabolic state of a cell. Therefore, we categorized the different amino acids into seven groups based on their metabolic origins (obtained from KEGG^17^), as shown in Fig 1B. These groups are derived from – glycolysis (ser, gly, ala), the pentose phosphate pathway PPP (phe, tyr, trp), from pyruvate (leu, ile, val), the alpha-ketoglutarate derived (glu, gln, asp, asn, thr), the glutamate derived (arg, pro, lys), sulfur amino acids (cys, met), and histidine (Fig 1B). Additionally, glutamate (and to a lesser extent glutamine) functions through a transamination reaction as an amine donor for other amino acids (Fig 1C), thus becoming a metabolic source for other amino acids. We utilize this grouping henceforth for its simplicity, representing the biosynthesis network in a glucose and nitrogen-replete environment.

**Figure 1:**
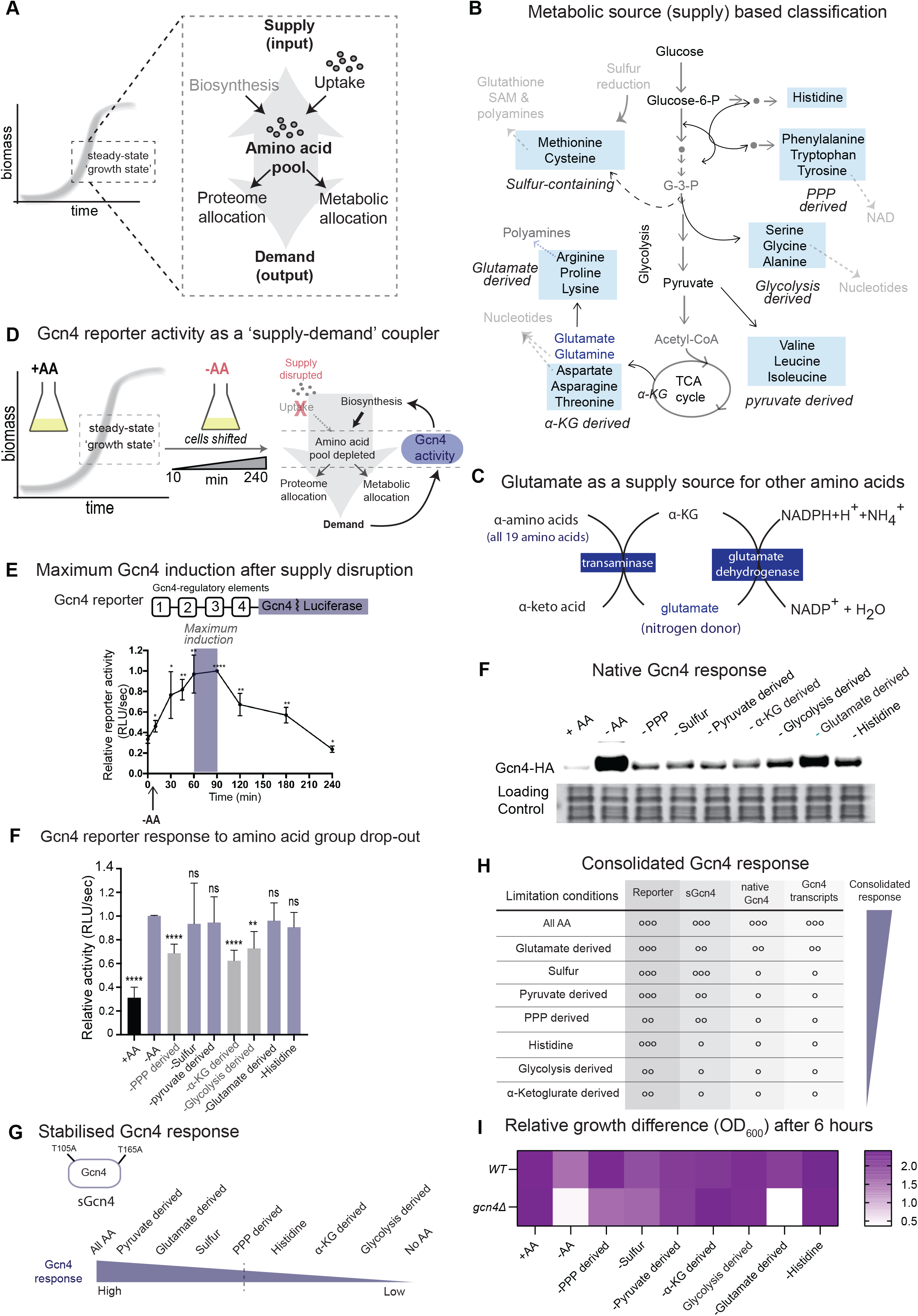
Cells exhibit hierarchical responses to supply-disruptions of amino acid groups, notably the glutamate-derived group. (A) Amino acid supply sources, and demand requirements of a growing cell. Supply includes amino acid biosynthesis and uptake from the environment. Demand includes the utilization of amino acids for translational and metabolic processes. (B) A metabolic source-based classification of all amino acids, forming seven groups derived from; glycolysis (ser, gly, ala), PPP (phe, tyr, trp), pyruvate-(leu, ile, val), alpha-ketoglutarate-(glu, gln, asp, asn, thr), glutamate-(arg, pro, lys), sulfur-(cys, met), and histidine. The figure highlights where in the central carbon network that each amino acid is derived from. (C) Glutamate (and glutamine) derived from the transamination of alpha-ketoglutarate, and as a metabolic source for all other amino acids. (D) Schematic illustrating how a transient disruption of amino acid supply can be achieved in batch culture, and how this supply-demand mismatch that can be perceived, managed and restored by Gcn4. (E) Experimental set-up and reporter for estimating responses upon transiently disrupting amino acid supply. The upper inset illustrates the *Gcn4-luciferase* reporter (26, 72) used. Cells were grown in +AA medium, harvested (time 0), washed with minimal medium (-AA), shifted to -AA medium, collected at the indicated time-points (10 to 240 min), and reporter activity measured. Relative luciferase activity is plotted, where activity at 90 min was set to 1. Data are displayed as means ± SD, n = 3. *p<0.05, **p<0.01, ***p<0.001, ****p<0.0001. (F) Assessment of reporter activity to the drop out of individual amino acid groups. Cells with the reporter were grown in +AA, washed with minimal media (-AA), shifted to different amino acid group dropouts for 75 min and relative reporter activity estimated. Activity in –AA medium was set to 1. Data are displayed as means ± SD, n = 4. *p<0.05, **p<0.01, ***p<0.001, ****p<0.0001, ns denotes non-significant difference. (G) Summary of stabilized Gcn4 (sGcn4) responses to the drop out of individual amino acid groups. Cells expressing sGcn4 (26) were grown in +AA, washed with minimal media (-AA), shifted to different amino acid group dropouts for 75 min and sGcn4 levels were estimated (see Fig S1b). The response observed is presented in the order of highest to lowest. (H) Native Gcn4 protein amounts in different amino acid dropout mediums, as detected using anti-HA antibody. A representative blot obtained from three biological replicates (n = 3) is shown. Also see Fig S1C, where relative band density in arbitrary units (A.U.) is plotted. *p<0.05, **p<0.01, and Fig S1D showing transcript abundance of direct Gcn4 targets, in the respective dropout conditions. (I) Summary of consolidated Gcn4 responses to the supply disruption of distinct amino acid groups. The scoring is based on the extent of Gcn4 response of the reporter, sGcn4, native Gcn4 and Gcn4 dependant transcripts in different amino acid dropouts, represented by the number of circles. The consolidated responses are arranged from highest to lowest. (J) Relative cell growth of wild-type and *11gcn4* cells in different amino acid group dropout conditions after 6 hours. Also see Figure S1D for quantifications.

With this starting point, we wanted to experimentally assess responses to transient disruptions in the supply of distinct amino acids. For this, we require a reliable reporter of mismatches in amino acid supply vs demand. Prototrophic yeast cells synthesize all amino acids when provided with nitrogen and carbon precursors, and would do so when external amino acid supply is broken. In order to restore supply during amino acid starvation, or maintain supply during growth, eukaryotic cells utilize the Gcn4/ATF4 transcription factor as an amino acid supply-hub to match demand^8, 13, 15^ (Fig 1D). If indeed cells differentially respond to transient disruptions in the supply of distinct amino acids, we hypothesized that the Gcn4 activity would correspondingly reflect this response. To investigate this, we shifted yeast cells growing in amino acid-replete medium (+AA), to a defined medium with no free amino acids (-AA), but with ammonium sulfate and glucose (to allow *de novo* amino acid biosynthesis). This would disrupt the extrinsic supply of all amino acids, requiring cells to restore supply via amino acid biosynthesis. We monitored the kinetics of Gcn4-activation using an established *Gcn4-luciferase* reporter^21, 22^, as shown in Fig 1D. Notably, the reporter activity increases within ∼15 minutes of shifting cells to -AA, and peaks at 60-90 min, after which it decreases (Fig 1E). This 60-90 min window of maximal activity (Fig 1D(i)) established a precise time-frame for subsequent analyses and is used henceforth. The kinetics of this observed response would also be consistent with that of a supply-demand coupler, which should increase in activity when supply is below demand, and then decrease as supply matches demand.

We next interrogated how this reporter activity changes when the supply of distinct amino acid groups are disrupted. Cells were grown in +AA medium and shifted to a synthetic medium where each amino acid group was dropped-out, but the other amino acids were supplemented (otherwise performed identically to experiments related to Fig 1E). Cells were collected at 75 minutes post-shift, and reporter activity estimated. The dropout of any amino acid group induced reporter activity to levels higher than amino acid replete medium (+AA) (Fig 1F), establishing that Gcn4 reports on supply-demand mismatches for all amino acids. However, the extent of reporter induction varied substantially for different amino acid groups (Fig 1F). Notably, the dropouts of sulfur (methionine and cysteine), pyruvate-derived (leucine, isoleucine and valine), glutamate-derived amino acids (arginine, proline and lysine), and histidine all showed robust reporter activity, comparable to complete amino acid drop out medium (-AA) (Fig 1F). In contrast, the dropouts of PPP-derived (tyrosine, tryptophan and phenylalanine), alpha-KG-derived (aspartate, asparagine, glutamate, glutamine and threonine), and glycolysis-derived amino acids (alanine, serine and glycine) showed significantly lower reporter activity (Fig 1F). This suggests possible prioritizations of restoration responses for distinct amino acid groups.

Gcn4 has sophisticated, multi-level post-transcriptional regulation^13, 23, 24^, as would be expected for a supply-controller. The final Gcn4 output comes from a combination of the translation of upstream regulatory elements of the Gcn4 transcript (as observed with the reporter), and the stability of the Gcn4 protein. We therefore further dissected the extent of Gcn4 regulation upon limiting the supply of different amino acid groups, by first assessing protein amounts of a stabilized Gcn4 protein (sGcn4-T105A, T165A)^21^, and native Gcn4. We observed higher sGcn4 protein in dropouts of sulfur-, pyruvate- and glutamate-derived amino acids (Fig S1B; lanes 5, 6, 9 and 1G). The dropout of PPP-derived amino acids also showed higher sGcn4 protein levels (Fig S1B; lane 4), minimal increase in sGcn4 in the histidine dropout (Fig S1B; lane 10). These results refined our earlier observations from the reporter, where sGcn4 strongly responds to the disruptions in sulfur, pyruvate-, glutamate-, and PPP- derived amino acids (Fig 1G). We next examined native Gcn4 protein after distinct amino acid group limitations. Native Gcn4 response was highest after the limitation of the glutamate-derived amino acids (Fig 1H; lane 8 and S1C). Finally, we estimated amounts of direct Gcn4- transcriptional outputs, as an end-point readout for this response. The transcripts of multiple, direct targets as obtained from^14^ increased the most in dropout medium for (respectively) sulfur-, pyruvate- and glutamate-derived amino acids, as well as PPP-derived amino acids (Fig S1D, with controls for Gcn4 RNA levels shown in Fig S1E). Through these multiple readouts (reporter, sGcn4, native Gcn4 and direct Gcn4-dependent transcripts) we construct a consolidated, hierarchically-graded Gcn4 response to supply-disruptions of distinct amino acid groups (Fig 1I). Cumulatively, the strongest response is to supply-disruptions of glutamate-derived amino acids, followed by strong responses to the sulfur-, and pyruvate-derived amino acids, a moderate response to PPP-derived amino acids, and a minimal response to the Glycolysis derived and alpha-KG derived amino acids in this condition (Fig 1I).

In complementary experiments, we examined the effect on short-term growth in cells lacking *GCN4* (*Δgcn4*), when each amino acid group was dropped-out. For this, the growth of wild-type and *Δgcn4* cells in different amino acid group dropout conditions were monitored. In these cells, the greatest relative reduction in growth was observed in the dropout of the glutamate-derived amino acids (Fig 1J and S1F). Together, these data reveal that upon transient amino acid supply disruption, cells are most constrained by the supply of glutamate-derived amino acids.

### Degree of response correlates with low biosynthetic cost and high total demand

We wondered how much of the observed responses reflected economic principles of supply and demand. To begin assessing this, we require estimates of individual amino acid supply costs, as well as how much total demand exists for distinct amino acids. We first calculated the cost to supply an individual molecule of each amino acid. For this we wanted to include as many components of cost as possible. Therefore, we accounted for all chemical reactions in each amino acid biosynthetic pathway (with glucose as the carbon source), and then computed the total high-energy phosphate bonds associated with each amino acid group, by including the amino acid precursors, metabolic precursors and reducing equivalents (NAD(P)H) required. Note: the complete calculations of metabolic are in Appendix 1, which includes a list of precursors used for each amino acid in the metabolic cost calculation in Table 1. The consolidated cost of all amino acid groups collectively comes from the *net* total number of high energy phosphate (ATP) produced (with NADH utilization), concurrent with total NADPH consumed (Appendix 1 and Table 2). The results of these cost estimations are shown in Fig 2A, as a relative-cost heat-map. From this, we find that for any single molecule of an amino acid, the glutamate-derived amino acids have the lowest supply costs. From these cost estimations, we also find that the highest response (which were observed for this group drop-out) correlated with low amino acid supply costs. To dissect this further, we assessed the individual costs of each of the glutamate-derived group - arginine, lysine and proline, wondering how different they were. Based on these cost estimations (in Fig 2A), arginine has the lowest supply costs, lysine has intermediate costs, and proline has very high costs. (Fig 2B, Appendix 1 and Table 1).

**Figure 2:**
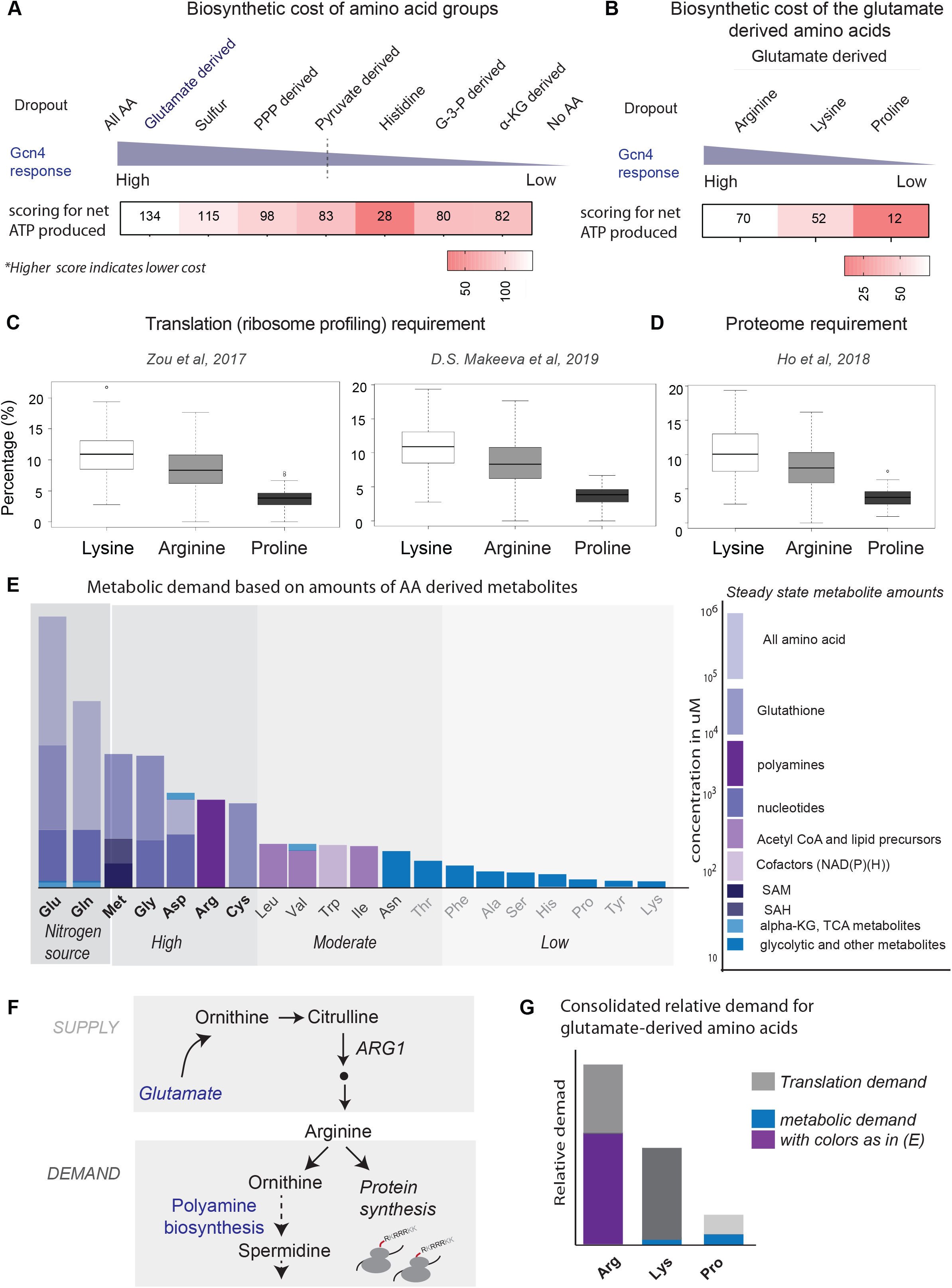
Degree of response correlates with low biosynthetic cost and high total demand. (A) Summary of the consolidated biosynthetic costs for distinct amino acid groups, with the grouping as in Fig 1. The consolidated biosynthetic cost comes from the total NADPH molecules consumed and total (net) number of ATP molecules *produced*, during the biosynthesis of the amino acids, and is for a glucose and ammonium replete condition. For complete details of all the calculations please see Appendix 1 and Table S2. (B) Summary of consolidated biosynthetic costs for each glutamate derived amino acid (arginine, lysine and proline) in a glucose and ammonium-replete condition. (C) Protein-synthesis demand for lysine, arginine and proline amino acids. Box plots showing the percentages of lysine, arginine and proline residues in the highly translated mRNAs, as assessed from two ribo-seq analyses datasets, (30) (left panel) and (31) (right panel). (D) Proteome demand for lysine, arginine and proline amino acids. Box plot showing the percentages of lysine, arginine and proline residues in the highly abundant proteins (primary dataset from (34)). (E) Qualitative estimates of demand for distinct amino acids. The different amino acids are organised in descending order based on their demands for metabolic outputs derived from each of these amino acids. The right-hand key indicates the intracellular concentrations of the different metabolic outputs derived from amino acids in the range of 100 of mMs to 100 of mMs. Different metabolic outputs are represented in different shades of blue. The size of the bars is scalable to the relative concentration of the metabolite in glucose and ammonium-replete condition (Higher concentrations reflected by taller bars). On the left, there is a graphical presentation for an ordering of amino acids based on high demand, moderate demand and low demand amino acids (assuming high glucose and ammonium). (F) Schematic illustrating the steps of arginine biosynthesis from Glutamate and the demands for arginine in the cell, which come from both protein synthesis, as well as for the production of polyamines. (G) Summary of the consolidated demand for each of the glutamate derived amino acids (arg, lys, pro), with a proportional relative-scale, included. Right hand key -The grey color scale (dark to light) representing the proteome/translation demand, high demand is represented in the darker shade than low demand. The blue color bars are the metabolic demand for the respective amino acid, which is same as (E). Arginine has high metabolic demand coming from polyamine biosynthesis. The size of the bar is scaled to the extent of demand.

**Table 1:**
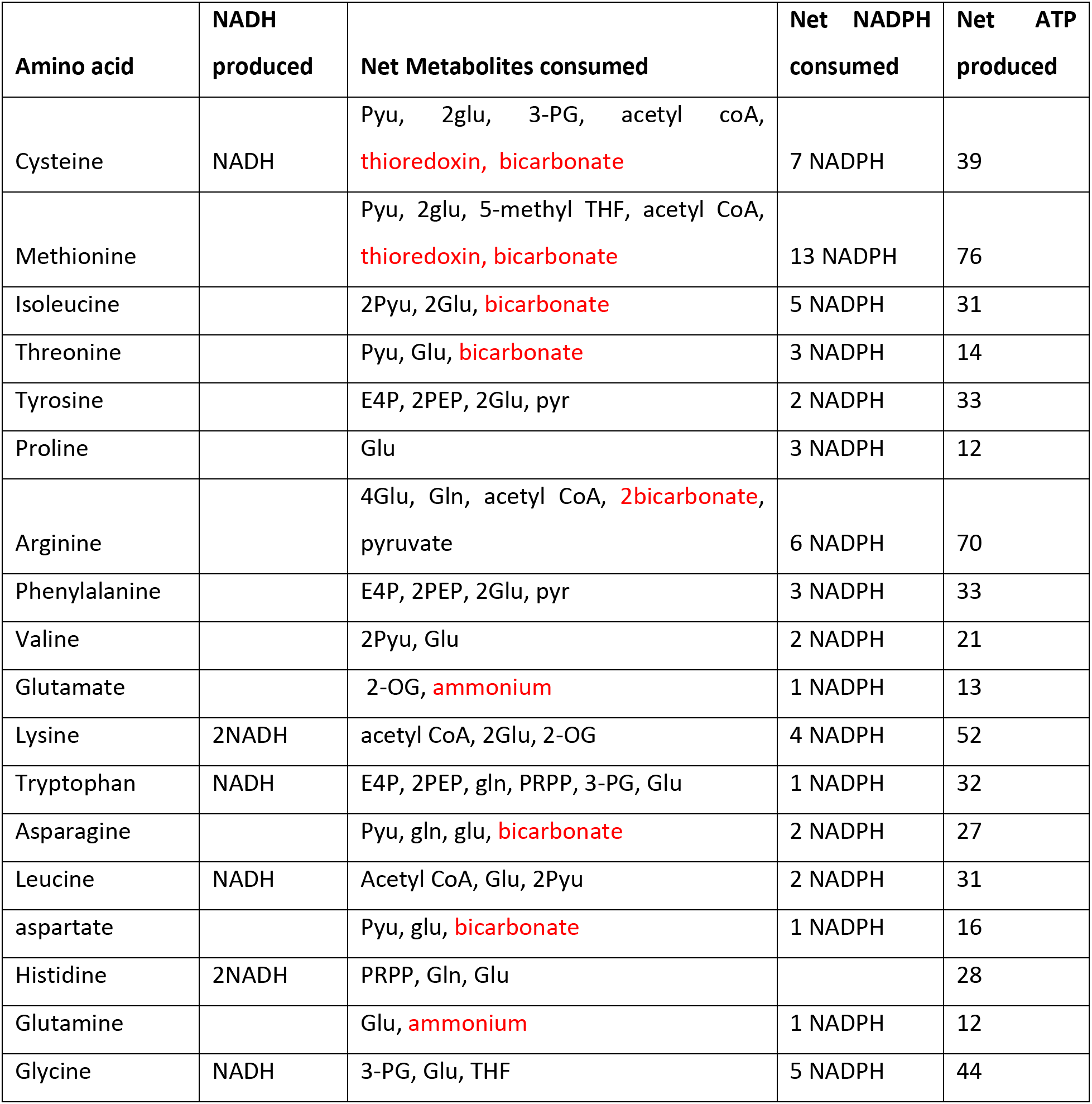
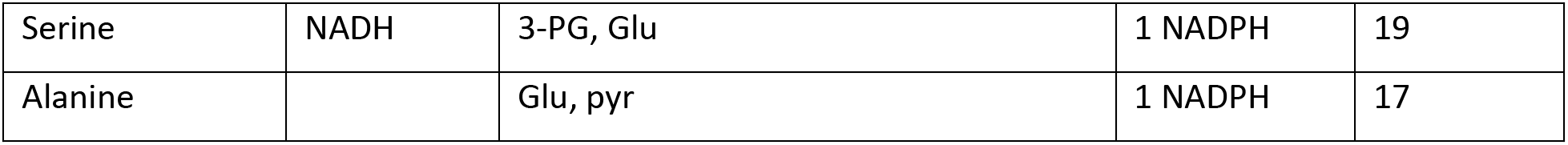
List of the metabolites consumed and produced by the cell to supply an individual molecule of each amino acid. The total number of NADH molecules produced, metabolites consumed, NADPH molecules consumed and ATP molecules produced for all 20 amino acids are shown. To calculate the number of NADPH molecules consumed and ATP molecules produced, all the chemical reactions in the biosynthetic pathway of all the metabolic precursors, except those shown in red, were accounted for, in cells growing with glucose as the carbon source. 1 NADH is equivalent to 3 ATPs.

**Table 2.** List of the metabolites consumed and produced by the cell to supply all the amino acids categorized in different groups.

| **Amino acid groups** | **Net ATP produced** | **Net NADPH consumed** |
| --- | --- | --- |
| sulfur | 115 | 20 |
| BCAA | 83 | 9 |
| TCA derived | 82 | 8 |
| Aromatic | 98 | 6 |
| Glutamate derived | 134 | 13 |
| Glycolysis derived | 80 | 7 |
| Histidine | 28 | 0 |

Having completed supply-cost assessments, we next asked what the composite demand requirement for distinct amino acids were. Currently, there are no estimates of total demand. Protein synthesis, which can be ∼quantitatively assessed, is one major demand for amino acids. For estimating demand coming from protein synthesis, we analyzed two ribosome-profiling datasets^25, 26^ (GEO accession: GSE91068 and GSE122039) from cells grown in comparable defined-medium conditions. The top 500 most highly translated genes from these datasets were grouped using gene ontology (GO). Expectedly, the most enriched GO category was the ribosomal constituents, consisting of ∼25% of the query genes (Fig S2A). This is consistent with our understanding that a majority of proteins in proliferating cells are ribosomal^27^. Next, we calculated the percentages of lysine, arginine and proline residues in these highly translated genes. Lysine, arginine and proline were ∼11%, 8.5% and 3.7% respectively in both the datasets (Fig 2C), and (as described earlier (13)), arginine and lysine are substantially overrepresented in the ribosomal proteins which form the translation machinery. We separately analyzed a whole-proteome dataset^28^, selected the top 500 most abundant proteins and performed a similar analysis as 2C. Again, the most enriched GO category was the ribosomal constituents, consisting of >15% of the query genes (Fig S2B). In this abundant protein set, lysine, arginine and proline were ∼10.4%, 8.1% and 3.8% respectively (Fig 2D), consistent with the ribo-seq analysis in Fig 2C. Together, these results show that the overall (high) demands from protein synthesis for lysine and arginine are comparable, and are ∼2-3 times greater than proline requirements.

This analysis of demand excluded the metabolic demand component for amino acids. We therefore built semi-qualitative but composite estimates for the demand for each amino acid, by adding up the ∼amounts of the primary metabolic outputs of every amino acid (Fig 2E, Table S3) (also see^29^). Given the extent of available data^29^, some estimates are order of magnitude based. We organized amino acids from highest to lowest metabolic demand (Fig 2E), and distinct amino acids fall into clear groups of high, moderate and low demand. This revealed a high metabolic demand for arginine, and low metabolic demands for lysine and proline (Fig 2E). Arginine is the largest assimilator of nitrogen in cells^30^, and much of the demand for arginine comes from its requirement in making polyamines (Fig 2E, 2F). Polyamines are abundant cellular metabolites, present in ∼millimolar amounts^31, 32^. In contrast, while the demand from protein synthesis for lysine is high, the metabolic demand for lysine is very low except in atypical contexts^33^. Finally, the demand for proline in protein synthesis is low, and there is no major metabolic demand for proline under the conditions used (Fig 2C, 2D, 2E). Therefore, for the group of glutamate-derived amino acids, the cumulative demand has a major contribution from the high arginine demand, a smaller contribution from the total lysine demand and the smallest contribution from the low demand for proline (Fig 2G). Summarizing our observations, arginine had the lowest biosynthetic costs of the glutamate-derived amino acids, but the highest demand.

### Disrupting arginine supply invokes the strongest response consistent with the law of demand

The magnitude of the restoration response seemed to correlate with high demand coupled with lower supply costs. In a given nutrient environment, supply parameters are inherent to the physico-chemical properties of that molecule, while demand has some component coming from selection and inherent growth rates for that cell. If these are primary criteria involved, how might cells prioritize an order in which to make amino acids when there are shortfalls? When supply parameters are inherent, the law of demand would drive the economy^34^. According to this law, the demand for an entity and its supply price are inversely correlated, and demand is highest for molecules with low supply costs^34^. In a growing cell, since amino acids amounts would be far from equilibrium^1, 4^, flux would be driven by the law of demand. If so, a hypothetical supply price vs demand curve for the glutamate-derived amino acids could be drawn as in Fig 3A.

**Figure 3:**
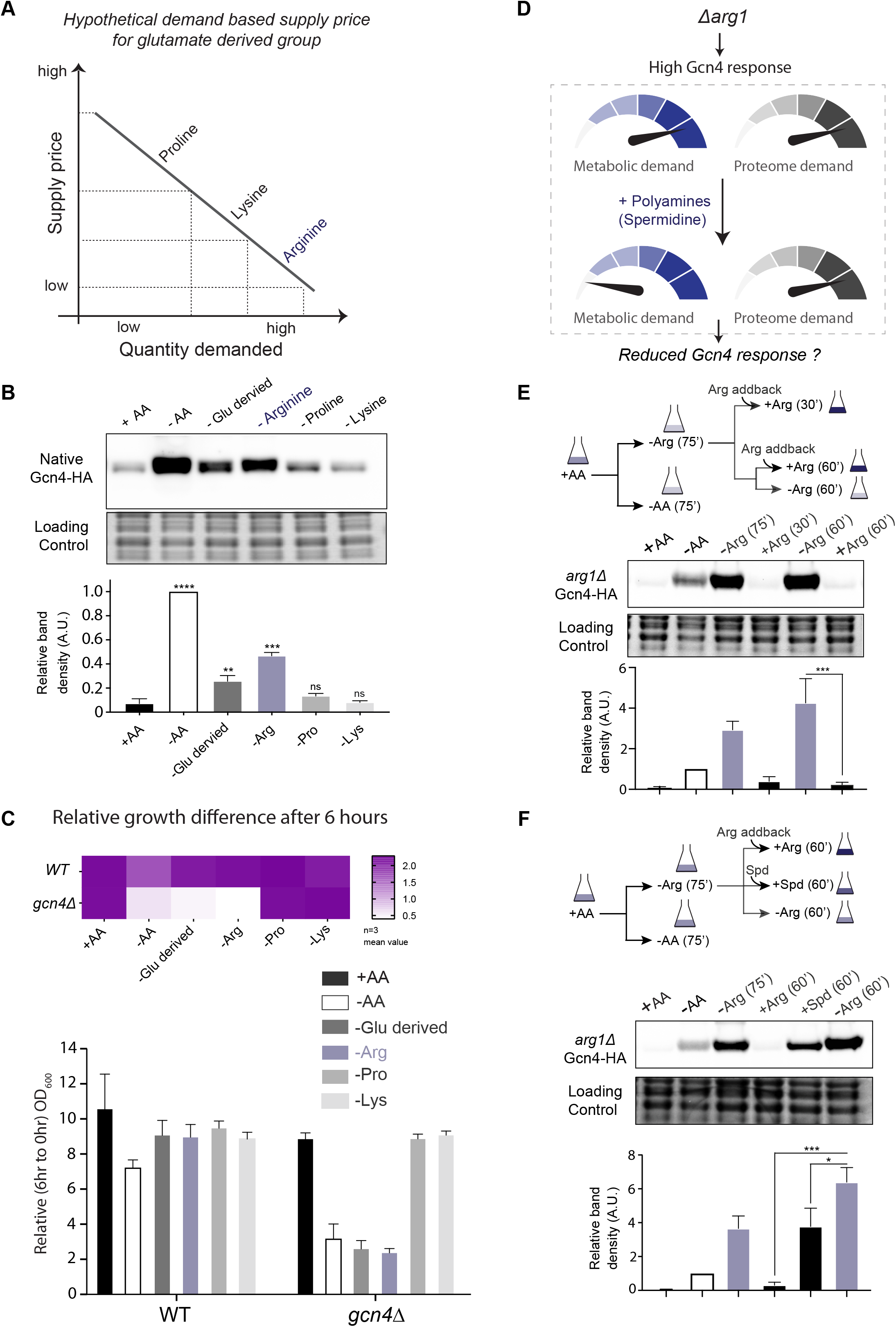
Arginine disruption invokes the strongest response consistent with the law of demand. (A) A hypothetical supply price vs required demand graph, as based on the law of demand, for the glutamate derived amino acids. (B) Gcn4 protein response for complete amino acid withdrawal (-AA), or for individual amino acid (arg, pro, lys) dropouts. A representative blot obtained from three biological replicates (n = 3) is shown. Relative protein amounts are shown below the representative blot, and comparisons are to the +AA condition. **p<0.01, ***p<0.001, ****p<0.0001, ns denotes non-significant difference. (C) Relative difference in the growth of wild-type and *Δgcn4* cells in different amino acid group dropouts (estimated after 6 hours). Data are displayed as means ± SD, n = 3. (D) Summary of predicted Gcn4 responses in *arg1Δ*, that would come due to high metabolic and proteome demand. If the polyamine contribution to this demand is significant, upon restoring polyamines, a reduction in Gcn4 would be expected. (E) Extent of Gcn4 induction upon arginine deprivation, in the absence of arginine biosynthesis. The upper schematic illustrates the experimental set-up. *arg1Δ* cells were grown in +AA, washed with -AA medium, shifted to either -AA or arginine dropout (-Arg) medium for 75 min and harvested. To the remaining cells in -Arg, arginine (10 mM) was added back (+arg) and incubated for 30 and 60 min. In parallel, cells were also shifted to arginine dropout (as a control) at the 60 min addback time-point, and Gcn4 amounts estimated in the different conditions. A representative blot obtained from four biological replicates (n = 4) is shown, along with quantifications. ***p<0.001. (F) Contribution of polyamines to the arginine-demand. The upper schematic illustrates the experimental set-up. Cells were grown in +AA, and washed in -AA, shifted to either -AA or arginine dropout medium for 75 min, and harvested. To the remaining cells growing in arginine dropout, either arginine (+Arg) or the major polyamine spermidine (+Spd) was supplemented, and incubated for 60 min. In parallel, cells were shifted to arginine dropout medium (control) for 60 min and Gcn4 amounts estimated in the different conditions. A representative blot obtained from three biological replicates (n = 3) is shown, along with quantifications. *p<0.05, ***p<0.001.

To experimentally test this possibility, we estimated the Gcn4 response to individual dropouts of arginine, lysine or proline (Fig 3B). Gcn4 protein was highest in the arginine dropout, as compared to lysine or proline dropouts (Fig 3B). Additionally, we estimated transcripts of direct Gcn4 targets. The arginine dropout had the highest response, compared to lysine and proline dropouts (Fig S3B). Indeed, this response in arginine dropout was comparable to complete dropout (-AA) (Fig 3B). In complementary experiments, we examined 6-hour growth in wild-type and *Δgcn4* cells in individual amino acid dropouts (for arginine, lysine and proline). *Δgcn4* cells showed the strongest growth reduction in the arginine dropout (Fig 3C and S3C). Together, these results indicate that cells are most sensitive to arginine limitation, amongst the glutamate-derived amino acids.

How much of the observed response upon arginine limitation might come from metabolic vs other demands? To estimate this, we established a system to address the extent of demand coming from metabolic requirements (polyamines). We used (*Δarg1*) cells, which cannot synthesize arginine, and therefore should display a constitutively high Gcn4 response to arginine limitation. Carefully designed add-back experiments, along with the corresponding estimation of Gcn4 protein responses, would allow us to address the contribution of arginine demand from polyamines (Fig 3D). Using *Δarg1* cells we first established a time-course to define the arginine addback response in –Arginine dropout medium (Fig S3D). The addback of arginine for ∼30 minutes reduced Gcn4 protein to ∼basal levels (Fig S3D). Next, we grew *Δarg1* cells in -Arginine medium for 75 min, supplied arginine for 30 and 60 minutes, and harvested cells (using 60 min arginine dropout (-Arg) as a control) (Fig 3E). Both the 30 and 60 min arginine addback reduced Gcn4 protein levels, with the 60 min addback resulting in the greatest reduction as compared to the -Arginine control (Fig 3E). With this system established, in order to test the polyamine contribution to the arginine dependent response, we grew *Δarg1* cells in - arginine medium as earlier (Fig 3E and S3D) and shifted cells to either arginine, or in medium supplemented with a major polyamine, spermidine (Fig 3F). Supplementing spermidine resulted in a ∼40% reduction in Gcn4 compared to the -Arginine control (Fig 3F, lanes 5 and 6). Expectedly, the arginine addback resulted in a ∼complete reduction in the level of Gcn4 (Fig 3F, lanes 4 and 6). These data reveal that ∼half the arginine demand in cells comes from metabolic requirements. Collectively, these data indicate that restoration-prioritization responses in cells for arginine function in accordance with the law of demand.

### Demand responses from TORC1 activity correlate with the law of demand

Conversely, the activity of the growth controlling TORC1 would reflect amino acid demand based on the growth state of the cell (Fig 4A). If this demand driven response were universal, whenever the cell needs to readjust demand after a supply disruption, then TORC1 activity should reciprocally reduce. We therefore assessed the TORC1 response upon transient arginine limitation, using the same experimental set-up as earlier. For this, we examined the phosphorylation status of a classical TORC1 target, Sch9^35^, after disrupting the supply of the glutamate-derived amino acids. TORC1 inhibition by rapamycin (which results in dephosphorylated Sch9) was used as a control for maximally reduced TORC1 activity (Fig 4B). Notably, the largest relative reduction of TORC1 activity was observed post arginine limitation, as compared to the dropouts of the other glutamate-derived amino acids (lysine or proline) (Fig 4B). We also independently assessed another major TORC1 output, which is the induction of ribosomal transcripts^36, 37^. Correspondingly, the greatest decrease in ribosomal transcripts (which reflect reduced TORC1 activity) was to arginine dropout (Fig 4C). Collectively, TORC1 activity shows the greatest reduction upon arginine disruption, and the Gcn4 and TORC1 responses to arginine are complementary, per the law of demand.

**Figure 4:**
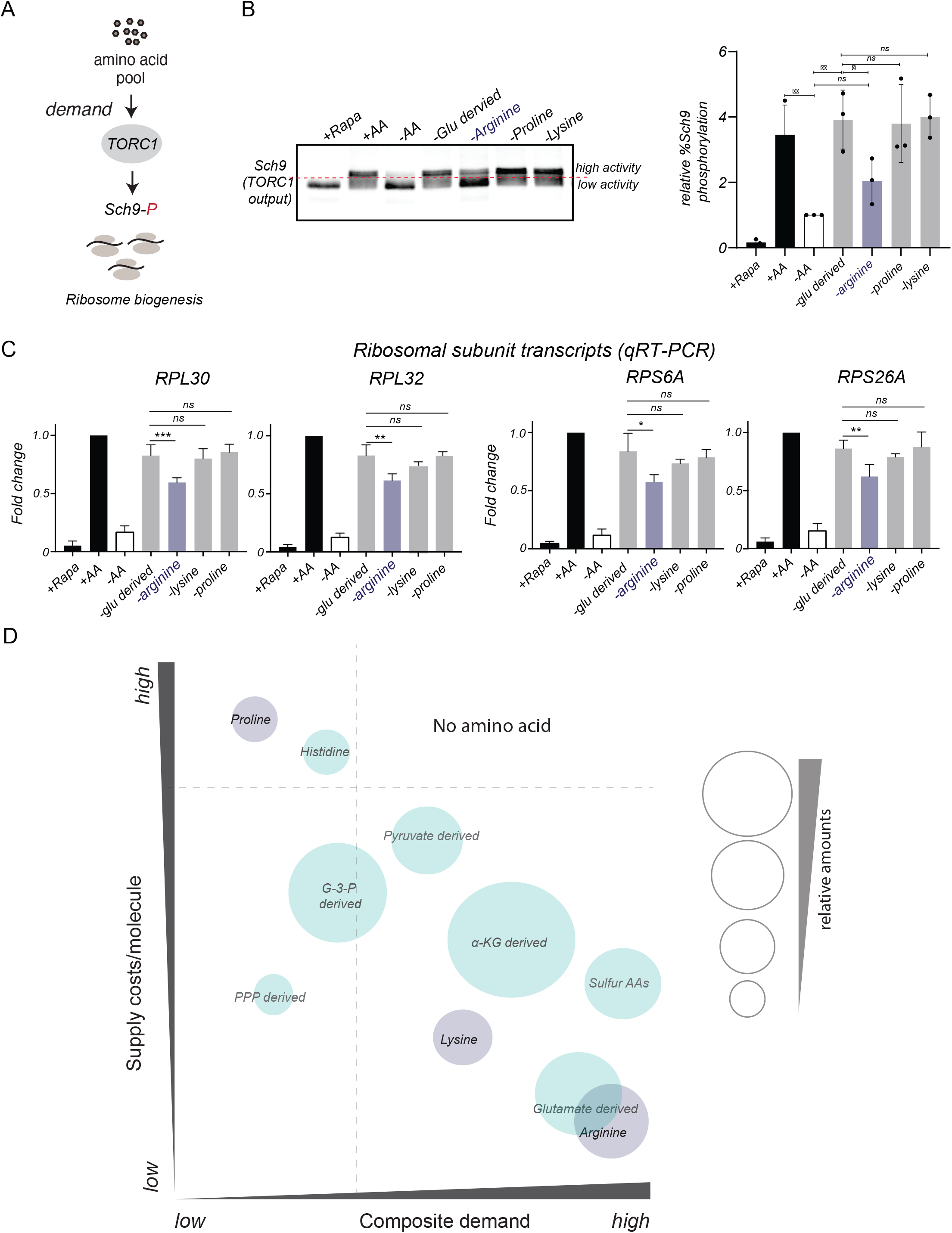
Demand based responses from TORC1 activity correlate with the law of demand. (A) Schematic illustrating the activity outputs of the TORC1, which indicates the extent of amino acid demand. Sch9 phosphorylation and ribosome biogenesis are key readouts of the extent of TORC1 activation. (B) Assessment of TORC1 activity upon transiently disrupting different amino acid supply (as indicated), based on Sch9 phosphorylation. Sch9 phosphorylation was assessed in different amino acid dropouts from the glutamate derived group (arg, pro, lys), using a classic assessment of electrophoretic mobility of Sch9 treated with NTCB (42), in different amino acid dropouts. A representative blot (from three biological replicates) is shown, and quantified (right-hand panel). Comparisons are to -AA conditions. ns denotes non-significant difference. **p<0.01, ***p<0.001, ****p<0.0001. (C) Assessment of TORC1 activity upon transiently disrupting different amino acid supply (as indicated), based on ribosomal transcript amounts. Relative changes in the expression of indicated ribosomal subunit transcripts, as TORC1-outputs, in different amino acid dropouts. Data are displayed as means ± SD, n = 3. ns denotes non-significant difference. **p<0.01, ***p<0.001, ****p<0.0001. (D) A model illustrating the relationship of the cost of supply for an individual molecule of an amino acid, with the total demand, in a four-quadrant chart. The size of the circles indicates (relative) intracellular concentrations of the respective amino acid in glucose and ammonium replete condition. Based on this framework, amino acids with low supply cost and low demand will elicit a lower response, and amino acids with low supply cost and high demand elicit the highest response. There are no amino acids which have both high supply costs as well as high demand.

## Discussion

In this study, we uncover hierarchical prioritization strategies to restore the supply of *distinct* amino acid groups, when supply is transiently disrupted. If this were organized based on both the supply costs along with demand, a model illustrated in Fig 4D emerges. Amino acids with low supply costs, but overall low demand in a given condition will elicit a minimal response because any supply disruption will be quickly restored. In this study’s context of glucose and nitrogen-replete environments, the glycolysis-derived amino acids have continuous and non-limiting precursor supplies, and fit these criteria. Although demand remains high for glutamate/glutamine, in the context of this study, the precursor supply is continuous and non-limiting (and they are essentially metabolic sources, as explained in Figure 1), resulting in a modest response. If individual molecule supply costs are high, but demand is low (such as proline), the limitation of such an amino acid will not matter much regardless of absolute costs to supply a molecule, since their supply restorations are easily managed. The strongest responses will therefore be for amino acids with modest individual supply costs, but high demand. This is because while demand is high, supply can still be restored. Such a scenario is exemplified by the response observed for arginine limitation. Based on this framework, there are no amino acids that have high supply costs and demand (Fig 4D). The prioritization strategies to restore the supply of amino acids, therefore, correspond to the law of demand.

While estimating supply costs for individual molecules was relatively straightforward, several assumptions were required in estimating allocations towards total demand that comes from a combination of metabolism and protein synthesis. Remarkable recent progress now allow excellent assessments of the resource allocations towards the proteome^6, 38, 39^. These however miss allocations of distinct amino acids towards metabolism. Contrastingly, systems-level metabolic output predictions do not require quantitative input data for all metabolites^40^, and many metabolic estimates are derived from ‘order of magnitude’ calculations^29^. By now including considerations of individual supply-costs and composite demand requirements, we can construct frameworks that suggest prioritization strategies towards distinct amino acids. We hypothesize the following, consistent with the law of demand. First, there are strict chemical constraints imposed in metabolism (based on available substrates, and the thermodynamics of reactions), resulting in finite outputs for amino acids^41^. Therefore, this component of amino acid allocations likely has stringent constraints and limited scope for large changes across organisms. Second, the allocations towards the proteome have greater scope for changes, based on the organism and the nature of its environment, growth and other selection. Consequently, across organisms, there can be large variations in proteome allocation strategies, and the cost estimates of supply and demand have to be made accordingly. This implicitly means that the model in Fig 4D will contextually change primarily in the lower two quadrants (of low costs, and high demand).

Our findings inform two directions of future inquiry, of a mechanistic nature. The first predicts what kinds of amino acid reserves might be needed. Intracellular concentrations of different amino acids vary extensively^18^, and growing cells maintain excess nitrogen reserves^6^. Including individual supply costs, as well as total demand could help (i) predict prioritization responses as the nutrient environment changes, and (ii) how much amino acid reserves different types of cells might need. An independent line of inquiry asks which amino acids might a cell need to sense? Amino acids are distinct based on chemical origins, and amounts required. By treating them as distinct entities based on these criteria, new and contextual roles for different amino acids in enabling cellular outcomes can be identified, coming from the demand side, or the supply side. On the demand side, while there are known *in vivo* TORC1 responses for specific amino acids^42–47^, many more responses are observed *in vitro* (63), suggesting undiscovered sensing systems. On the supply side, the Gcn2/Gcn4 axis itself can distinguish between individual amino acid groups and is also activated independent of amino acid starvation^48^.

Living organisms in Miller’s “Living Systems” have been compared to factories, with cells described as open systems, dealing with multiple inputs and outputs of matter, energy and information^49^. In this context, how much can prioritizations of resource restorations be understood based on demand-dependent criteria (or in other words, when would demand dominate)? For cells, a main resource constraint is the input price, which is governed by precursor availability, and rules of chemistry, thermodynamics and evolutionary history. For amino acids, the individual supply prices are fairly constant. Therefore, demand-based criteria would dominate in cells when concentrations of metabolites are saturating (above the enzyme km values), and this is typically the case for amino acids, resulting in a demand-elasticity of zero^1–3, 50^. In contrasting contexts, where metabolite consumptions are regulated as a function of available supply, supply-based criteria will become increasingly important^3^. Additionally, in a demand-driven context cells must manage the total costs of supply because if an input cost is high, the supply is limited since there are boundaries to how much energy can be obtained within cells. An inference from this is that cells would optimize their outputs in tune with these energetic constraints^51, 52^. From classical economics, other factors that determine supply prices are: (i) processes used for supply, (ii) anticipation of future prices, (iii) the number of suppliers, and (iv) the presence of monopolies or cartels that restrict supply or increase perceived value of goods^34^. In cells the first (enzymes, transporters) are optimized by selection, evidence for the second is limited to systems exhibiting hysteresis or oscillatory behavior, the third is finite and quantifiable, and the fourth rarely exists. Cells do not consider ‘Veblen goods’^53^, which have artificially high prices because of perceptions (eg. as status symbols) that manufacture demand. These considerations could therefore help identify resource bottlenecks, as well as likely responses to disruptions in specific resources, and distinguish supply-driven vs demand-driven economies. Work in these areas holds promise to advance a broader understanding of resource allocation strategies in cells and to improve the metabolic engineering of cell factories.

## Materials and Methods

### Yeast strains, media and growth conditions

The prototrophic CEN.PK strain of *Saccharomyces cerevisiae* was used as the wild type in all the experiments ^54^. All the strains used in this study are listed in Table S3. For all experiments in complete medium, an overnight preculture was grown at 30°C (to OD_600_ ∼2) in YPD medium (1% yeast extract, 2% peptone, 2% dextrose), and subsequently sub-cultured in the same media (starting OD_600_ 0.2) and grown to an OD_600_ ∼0.5-0.6. Rapamycin treatment: cells were treated with 200 nM rapamycin for 75 min and then harvested. For amino acid limitation experiments, the cells grown as above were washed once with defined **minimal medium** (0.67% yeast nitrogen base with ammonium sulfate, without amino acids, 2% dextrose) and shifted to this medium to which specified amino acid were supplemented. Cells were grown in this medium for 75 min and then collected. For strains bearing plasmid, appropriate selection was used for growing the overnight culture. The strains and plasmids used in this study are listed in Table S1 and S2 respectively.

### Defined and amino acid dropout medium

For reconstitution of different amino acid dropout media, all amino acids were added to a final concentration of 2 mM each except for cysteine which was added to a final concentration of 0.5 mM. Tyrosine was excluded from the medium due to its poor solubility. The amino acid dropout media used are listed in Table S3. For the addback experiments, 10mM arginine and spermidine was used.

### Luciferase reporter assay for Gcn4 translation

Luciferase assay was performed as described previously ^21, 22^. Briefly, wild-type cells transformed with a Gcn4-luciferase reporter were grown overnight in YPD with G418. Cells were sub-cultured in the same media without selection starting at OD_600_ of 0.2 and grown till the OD_600_ reached ∼0.5. 20 ml of this culture was collected (“0 min”) and pellets frozen at -80 ⁰C. The remaining culture was collected, washed and shifted to the specified minimal medium and incubated at 30 ⁰C for indicated time-points. For the kinetics experiments, cells were harvested after 15, 30, 45, 60, 90, 120, 180 and 240 min. Cell pellets were stored at -80 ⁰C. Pellets were resuspended and lysed in lysis buffer, cleared by centrifugation, and protein concentrations estimated. Luciferase assays were performed using luciferase assay system (E1500, Promega) and activity measured using a Sirius luminometer (Tiertek Berthold). Data was obtained as Relative Light Units per sec (RLU/s). Statistical significance was determined using a Student T-test (GraphPad Prism 7). For single-time point experiment, cells were shifted to the indicated minimal medium, cells harvested after 75 min and luciferase assays were performed as described previously.

### Addback experiments with arginine and spermidine

Cells were grown overnight and then sub-cultured in YPD as described earlier. Cells were then washed once with minimal media and then shifted to minimal medium or minimal medium with arginine dropout for 75 mins. At 75 minutes, 20 OD_600_ from the cultures were harvested. The remaining cells were then divided for the dropout/addback experiments. For addback experiments 10mM arginine or spermidine were added back at the indicated time point, and cells collected for further analysis.

### Western blot analysis

Cells were collected, proteins extracted and estimated, and protein amounts were estimated using Western blotting with specific antibodies, as described earlier^21^. All western blots were done in triplicates. For stabilized Gcn4 protein levels, the SL161 plasmid with stabilized Gcn4-HA (Table S3) was expressed in *gcn4*Δ strain, and analyzed as described earlier^21^. For steady-state Gcn4 protein levels, Gcn4 that is tagged at its chromosomally locus with a C-terminal HA epitope^21, 55^ was used. For Sch9 detection, Sch9 was tagged at its chromosomal locus with a HA epitope. 10-12 OD_600_ of these cells were collected with 6% final concentration of TCA, kept on ice for 15 min, harvested and washed twice with ice cold acetone by centrifugation at 3500 rpm, 3 min, 4°C. The cell pellets were dried in a speed-vac for 45-60 min and stored at -80°C. Cell pellets were resuspended in 300 µl of urea lysis buffer and processed for NTCB (2-Nitro-5-thiocyanatobenzoic acid) cleavage (to assess its phosphorylation status and TORC1 activity) as described previously^35^. Cleaved Sch9 was detected using anti-HA (H6908-2ML Sigma-Aldrich) primary antibody and anti-rabbit horseradish peroxidase-conjugated (7074S, Cell Signaling technologies) secondary antibody.

### RNA isolation and quantitative real-time PCR (qRT-PCR) analysis

Cells were grown in the specified media, and 5-6 OD_600_ of cells were harvested. RNA was isolated by a hot phenol beating method as described earlier^22^, treated with DNAseI (AM2238, Thermo Fisher), cDNA synthesized with random primers (48190011, Thermo Fisher) and SuperScript III reverse transcriptase (18080-085, Thermo Fisher Scientific) according to the manufacturer’s protocol. Relative transcript quantifications were done by real-time PCR on an ViiA 7 Real-Time PCR System (Thermo Fisher) using Maxima SYBR Green/ROX qPCR Master Mix (K0222, Thermo Fisher). *ACT1* was used as an internal normalization control. All qRT-PCRs were performed in triplicates using three independent biological RNA samples. Statistical significance was determined using a Student T-test (GraphPad Prism 7).

### Growth assays conditions

WT and *gcn4*Δ cells were grown in YPD and sub-cultured in fresh minimal, dropout or complete medium and cell growth monitored using O.D_600_ for a total of 11hrs. For final comparisons, relative growth at 0hr, 6hr and 11hr were compared.

## Acknowledgements

SL acknowledges DBT-inStem, and a DBT-Wellcome Trust India Alliance Senior fellowship (IA/S/21/2/505922) for support.

---

**Supplementary figure S1:**
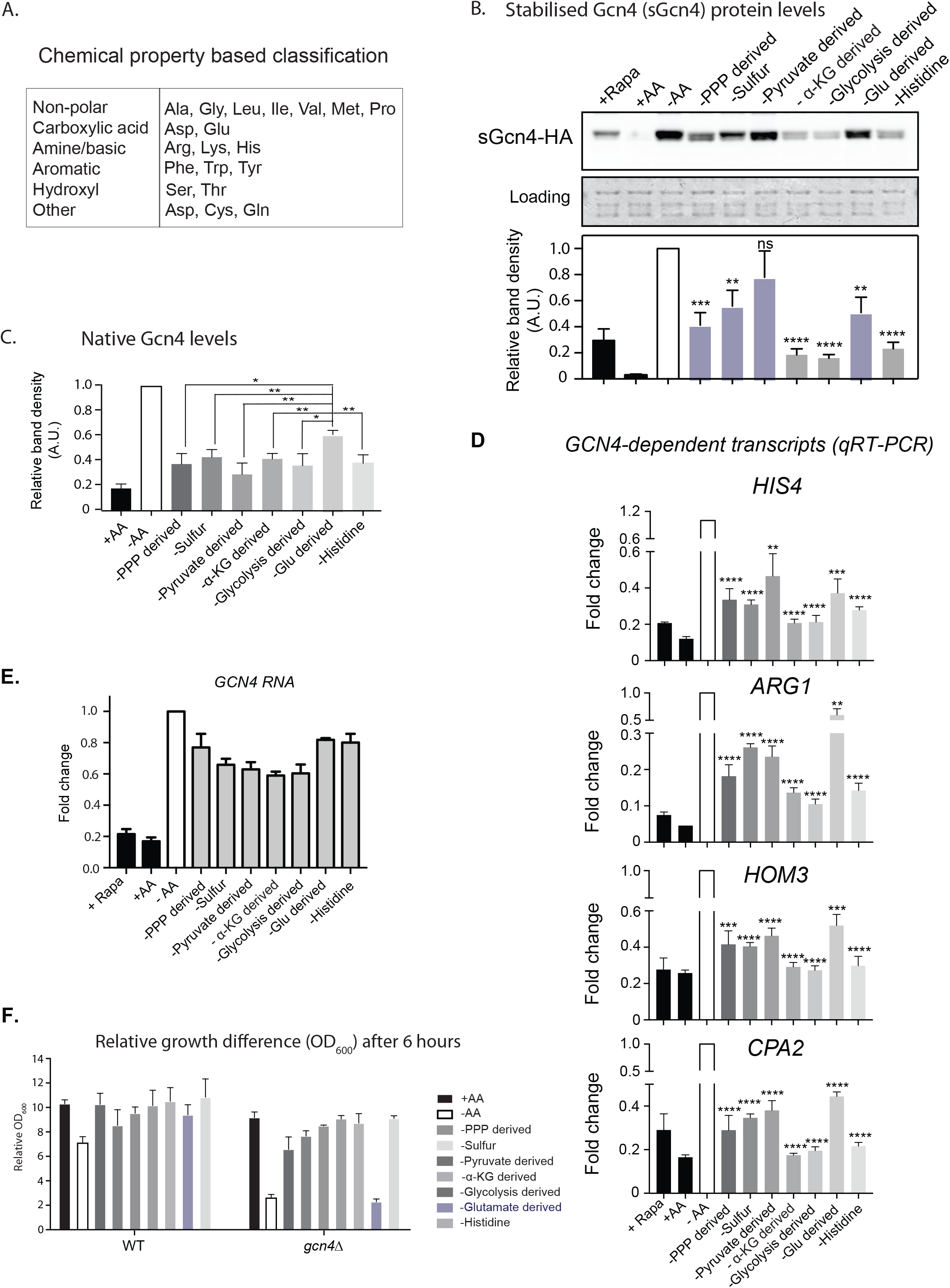
(A) A conventional (chemical property) based classification of amino acids^16^ (B) Stabilized (sGcn4) protein levels (*Δgcn4* cells transformed with a stabilized Gcn4 (T105A, T165A)) in different amino acid dropout media, detected using an anti-HA antibody. A representative blot obtained from three biological replicates (n = 3) is shown. **p<0.01, ***p<0.001, ****p<0.0001, ns denotes non-significant difference. (C) Relative intensity of native Gcn4 protein, from three biological replicates (representative blot in Fig1G) in arbitrary units is plotted. **p<0.05, ***p<0.01. (D) Transcript abundance of Gcn4 targets for n=3 in respective amino acid group dropouts. **p<0.01, ***p<0.001, ****p<0.0001, ns denotes non-significant difference. (E) Relative changes in the *GCN4* mRNA levels in respective dropouts of the amino acids. *GCN4* mRNA levels remained largely unchanged within the groups. (F) Relative biomass (cell growth) changes of WT and *flgcn4* after 6 hours of growth. The OD_600_ was normalised to 0 hour and compared (n=2).

**Supplementary figure S2:**
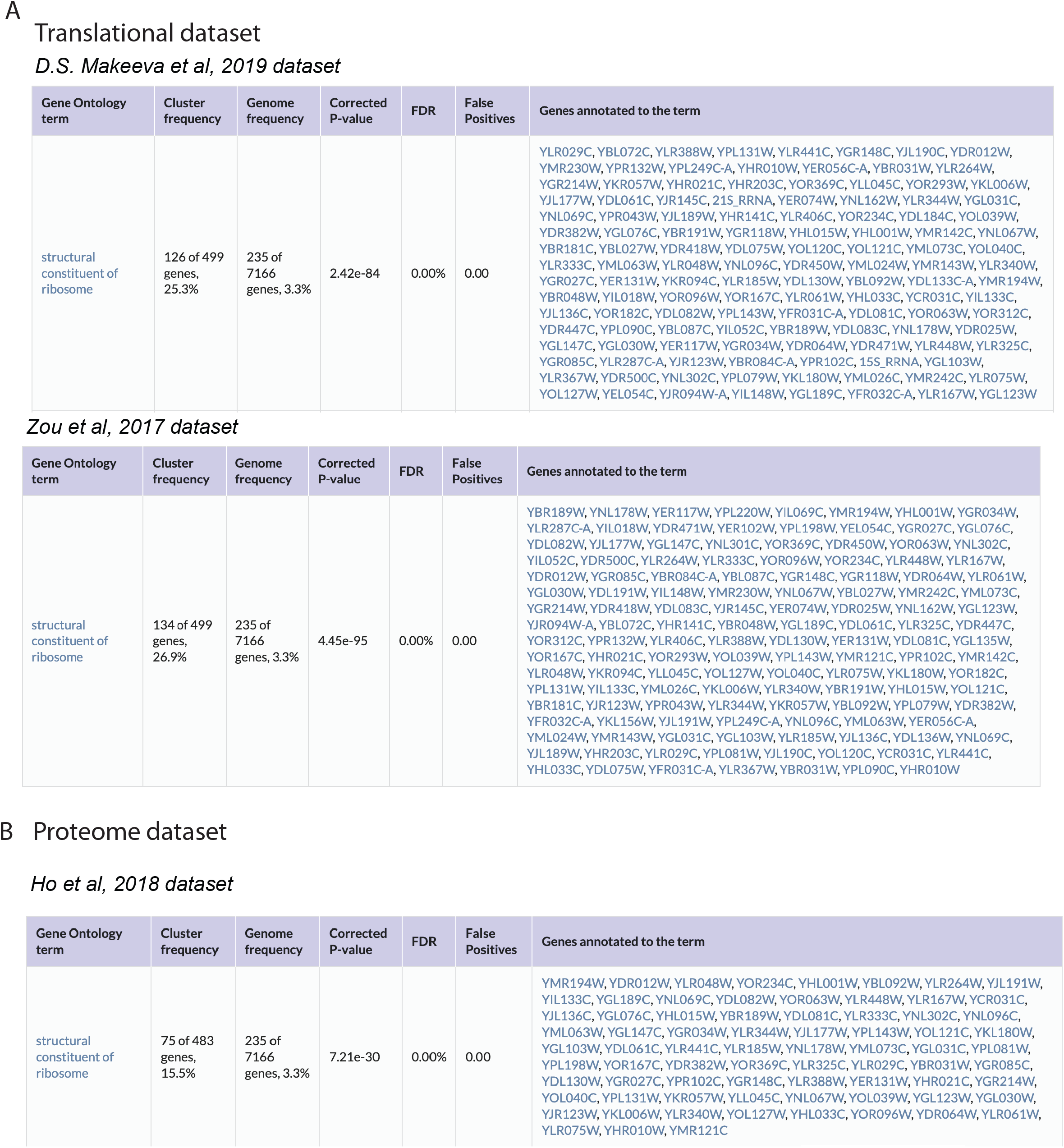
(A) The translational datasets used to estimate the translational demand of glutamate-derived amino acids in Fig2. (B) The proteome datasets used to estimate the translational demand of glutamate-derived amino acids in Fig2.

**Supplementary figure S3:**
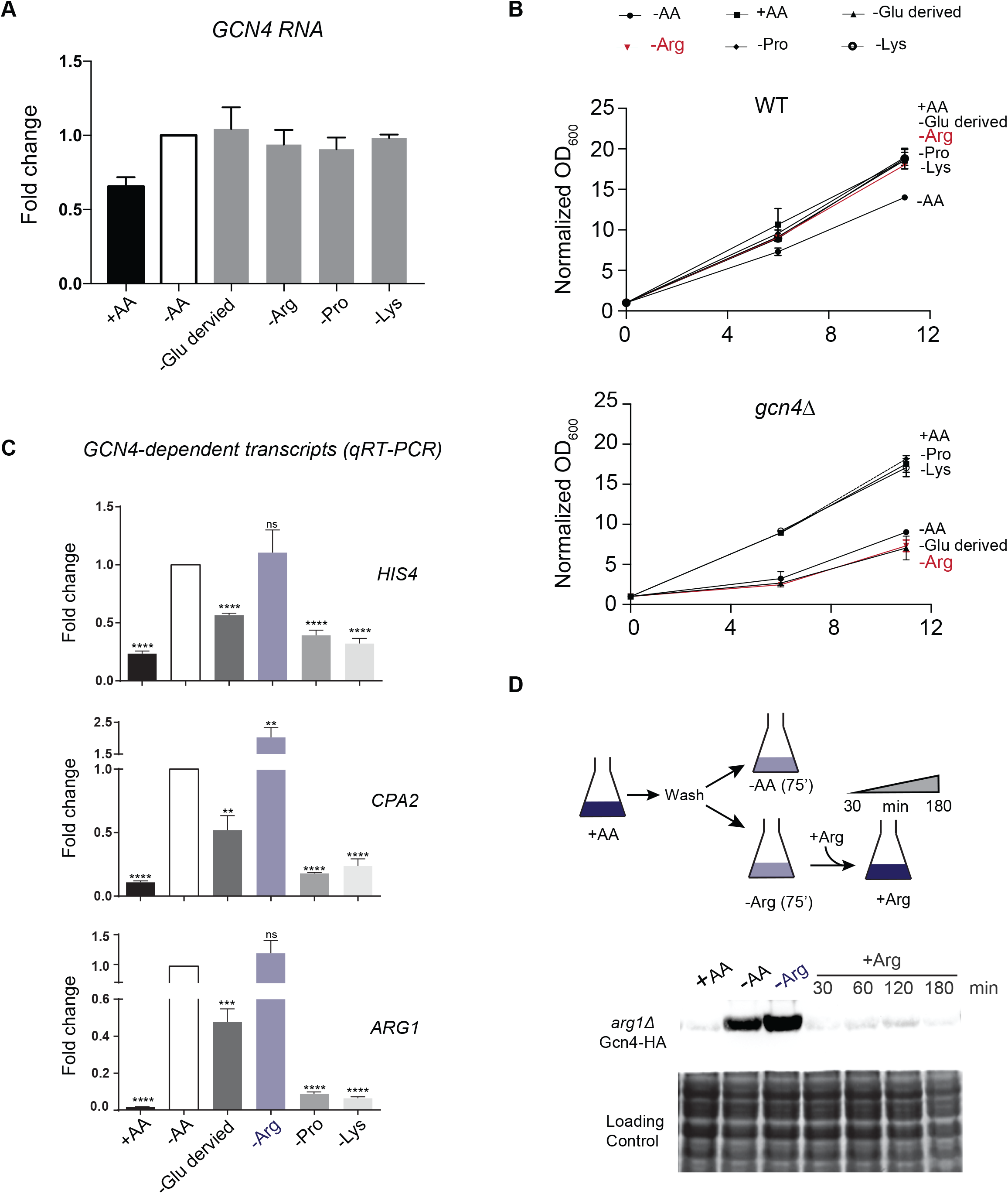
(A) Relative *GCN4* mRNA amounts in different amino acid dropout media (glutamate derived group). *GCN4* mRNA does not show any significant difference between all amino acid dropouts and other amino acid dropouts. (B) Relative growth of *Δgcn4* cells over 12 hours, in different drop out medium. *Δgcn4* cells showed the strongest growth reduction in the arginine drop out, and this was comparable to that of all glutamate-derived amino acid (arg+lys+pro) dropout combined. (C) Fold change in Gcn4 transcripts in Arg, Pro, Lys dropout media is plotted for n=3. **p<0.01, ***p<0.001, ****p<0.0001, ns denotes non-significant difference. (D) Schematic for time course experiment for the arginine addback to cells starved for arginine in *arg1Δ* cells. Western blot showing Gcn4 amounts in the indicated medium. The highest Gcn4 protein was in -arginine medium, and Gcn4 amounts reduced upon adding 10mM arginine to the medium at different time points.

## Supplementary Tables

**Table S1.** Strains used in this study.

Strains used in this study
| *Strain background* | *Genotype* | *Source* |
| --- | --- | --- |
| CEN.PK | *Mat a* | Van Dijken et al, 2000 |
| *gcn4Δ* | *Mat a gcn4Δ::NAT* | Walvekar et al 2018 |
| *GCN4-HA* | *Mat a GCN4-HA-KanMX* | Walvekar et al 2018,<br>Srinivasan et al 2020 |
| *SCH9-HA* | *Mat a SCH9-HA-NAT* | This study |
| *GCN4-Flag pdr5Δ* | *Mat a GCN4-flag-G418 pdr5Δ::NAT* | Walvekar et al 2018,<br>Srinivasan et al 2020 |
| *arg1Δ GCN4-HA* | *Mat a arg1Δ::NAT GCN4-HA-KanMX* | This study |

**Table S2.**
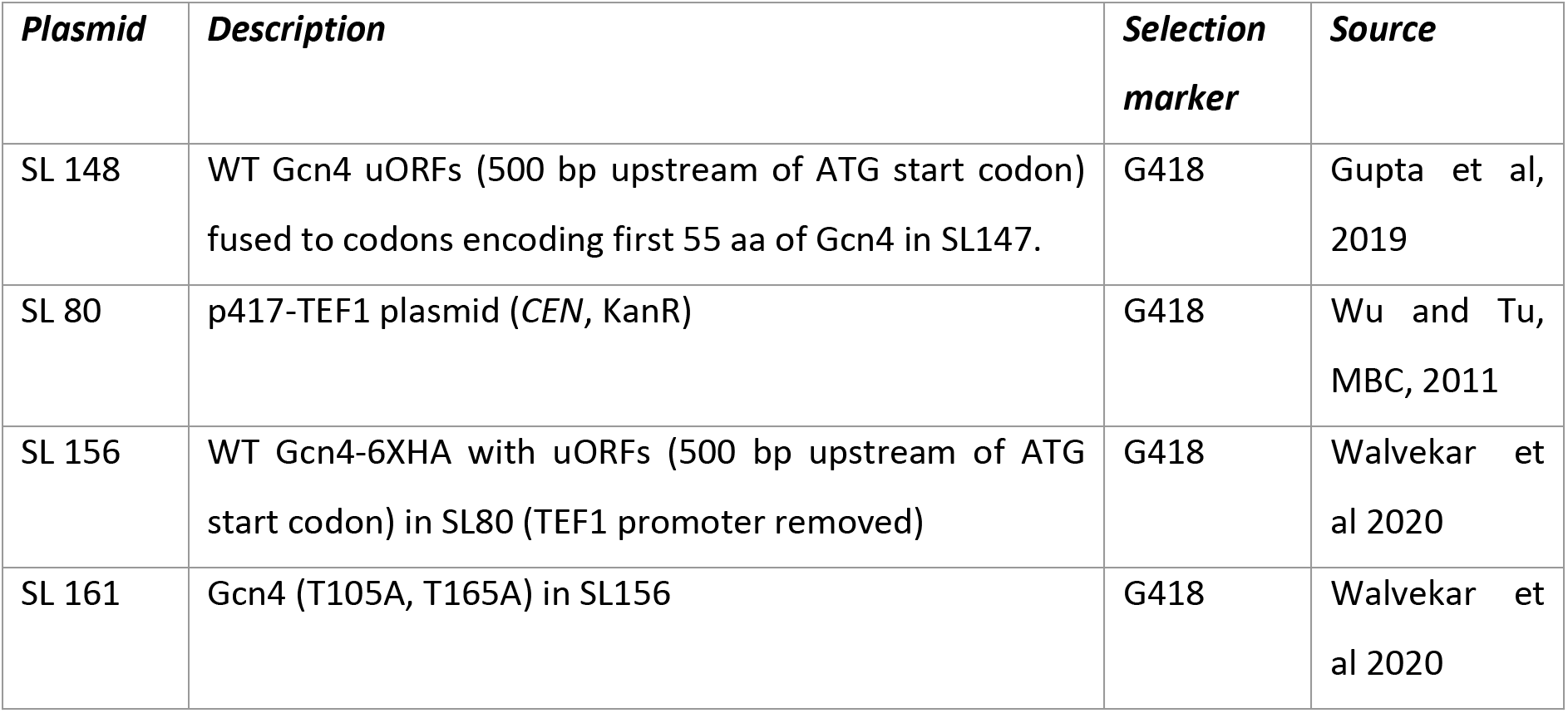
Plasmids used in this study.

Plasmids used in this study
| *Plasmid* | *Description* | *Selection marker* | *Source* |
| --- | --- | --- | --- |
| SL 148 | WT Gcn4 uORFs (500 bp upstream of ATG start codon) fused to codons encoding first 55 aa of Gcn4 in SL147. | G418 | Gupta et al, 2019 |
| SL 80 | p417-TEF1 plasmid (*CEN*, KanR) | G418 | Wu and Tu, MBC, 2011 |
| SL 156 | WT Gcn4-6XHA with uORFs (500 bp upstream of ATG start codon) in SL80 (TEF1 promoter removed) | G418 | Walvekar et al 2020 |
| SL 161 | Gcn4 (T105A, T165A) in SL156 | G418 | Walvekar et al 2020 |

**Table S3.**
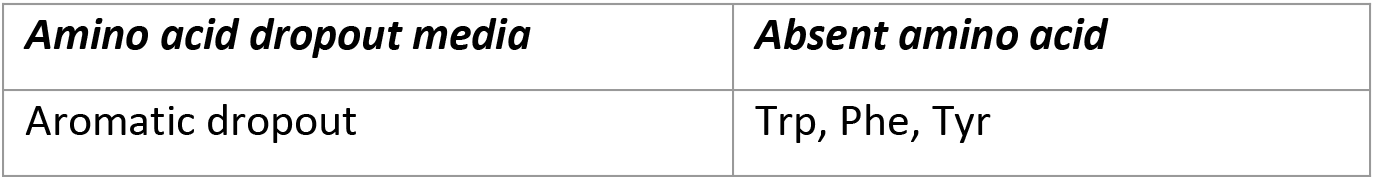
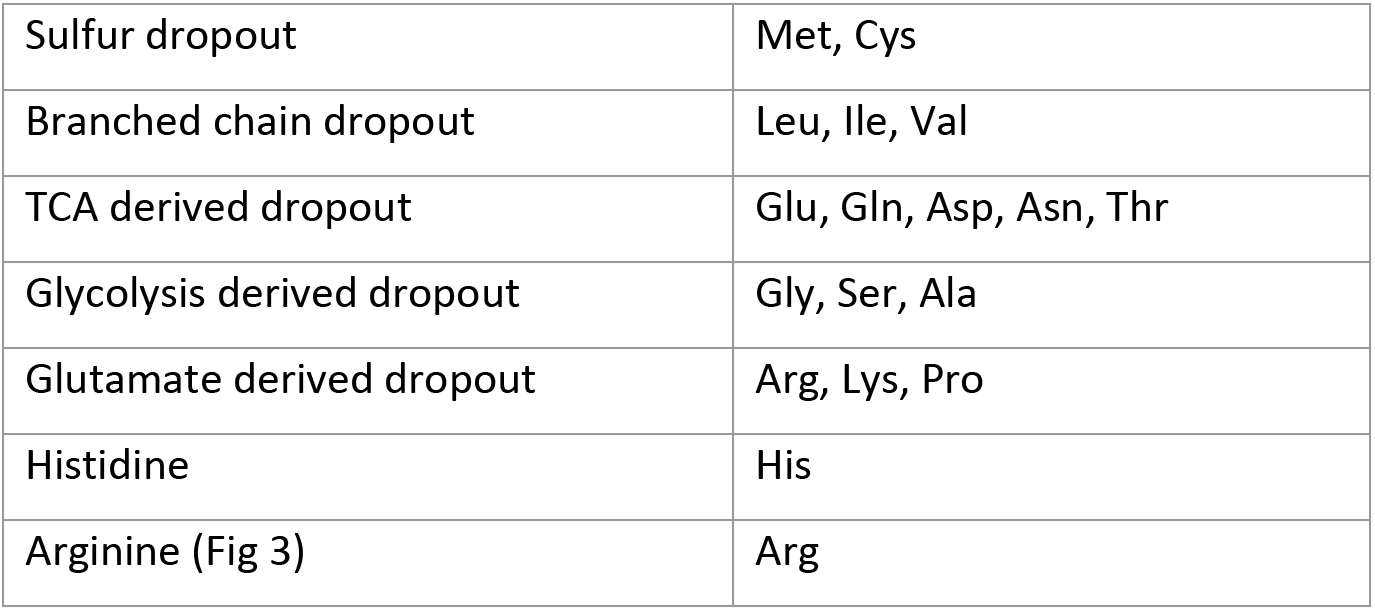
Amino acid dropout media used in this study.

Amino acid dropout media used in this study
| *Amino acid dropout media* | *Absent amino acid* |
| --- | --- |
| Aromatic dropout | Trp, Phe, Tyr |
| Sulfur dropout | Met, Cys |
| Branched chain dropout | Leu, Ile, Val |
| TCA derived dropout | Glu, Gln, Asp, Asn, Thr |
| Glycolysis derived dropout | Gly, Ser, Ala |
| Glutamate derived dropout | Arg, Lys, Pro |
| Histidine | His |
| Arginine (Fig 3) | Arg |

## Appendix 1 Extended Methods

### 1. Basis for calculation of biosynthetic cost of all amino acids in yeast

For the calculation of biosynthetic cost of each amino acid, we accounted for all the chemical reactions in the biosynthetic pathway of amino acids in cells growing with glucose as the carbon source. All amino acids are synthesized in the cell from either the intermediates of glycolytic pathway, pentose phosphate pathway (PPP) and tricarboxylic acid (TCA) cycle. This conversion of metabolic intermediates to an amino acid molecule involves several chemical reactions. Based on the nature of the chemical reactions, there is either energy consumption or production. The energetic cost is in the form of either high energy phosphate bonds (ATP) or reducing equivalents (NADH, NADPH). During oxidative phosphorylation, 3 ATP molecules are generated from 1 NADH molecule. Thus, the consolidated energetic cost for biosynthesis of each amino acid molecule is calculated by considering the number of the ATP molecules consumed/produced and by converting the number of the NADH molecules to ATP molecules using the conversion, 1 NADH =3 ATP molecules. In addition to this direct energetic cost, some chemical conversions also require other metabolic precursors or cofactors, which also have an associated biosynthetic cost.

#### Biosynthetic cost calculation for each amino acid

1. **Methionine**: 5 ATP and 5 NADPH molecules are consumed. Amino acid precursors consumed are: 2 glutamate (glu) and other metabolic precursors consumed are: 1 acetyl CoA (acCoA), 1 thioredoxin, 1 bicarbonate, 1 5-methyl tetrahydrofolate (5-THF) and 1 pyruvate.
2. **Cysteine**: 4 ATP and 5 NADPH molecules are consumed. Amino acid precursors consumed are: 2 glutamate (glu) and other metabolic precursors consumed are: 1 acetyl CoA (acCoA), 1 thioredoxin, 1 bicarbonate, 1 pyruvate and 1 3-phosphoglycerate (3-PG).
3. **Serine**: 1 NADH molecule is produced. Amino acid precursor consumed is: 1 glutamate (glu) and other metabolic precursor consumed is: 1 3-phosphoglycerate (3-PG).
4. **Alanine**: amino acid precursor consumed is: 1 glutamate (glu) and other metabolic precursor consumed is: 1 pyruvate.
5. **Glycine**: 1 NADH molecule is produced. Amino acid precursor consumed is: 1 glutamate (glu) and other metabolic precursors consumed are: 1 3-phosphoglycerate (3-PG) and 1 tetrahydrofolate (THF).
6. **Aspartate**: 1 ATP molecule is consumed. Amino acid precursor consumed is: 1 glutamate (glu) and other metabolic precursors consumed are: 1 pyruvate and 1 bicarbonate.
7. **Glutamine**: 1 ATP molecule is consumed. Amino acid precursor consumed is: 1 glutamate (glu).
8. **Glutamate**: 1 NADPH molecule is consumed. Metabolic precursor consumed is: 1 2-oxoglutaric acid (2-OG).
9. **Asparagine**: 2 ATP molecules are consumed. Amino acid precursors consumed are: 1 glutamate (glu), 1 glutamine (gln) and other metabolic precursors consumed are: 1 bicarbonate and 1 pyruvate.
10. **Threonine**: 3 ATP and 2 NADPH molecules are consumed. Amino acid precursor consumed is: 1 glutamate (glu) and other metabolic precursors consumed are: 1 bicarbonate and 1 pyruvate.
11. **Tryptophan**: 1 ATP and 1 NADPH molecules are consumed and 1 NADH is produced. Amino acid precursors consumed are: 1 glutamate (glu), 1 glutamine (gln) and other metabolic precursors consumed are: 1 5-phosphoribosyl pyrophosphate (PRPP), 1 3-phosphoglycerate (3-PG), 1 erythrose 4-phosphate (E4P) and 2 phosphoenolpyruvate (PEP).
12. **Tyrosine**: 1 ATP molecule is consumed. Amino acid precursors consumed are: 2 glutamate (glu) and other metabolic precursors consumed are: 1 pyruvate, 1 erythrose 4-phosphate (E4P) and 2 phosphoenolpyruvate (PEP).
13. **Phenylalanine**: 1 ATP and 1 NADPH molecules are consumed. Amino acid precursors consumed are: 2 glutamate (glu) and other metabolic precursors consumed are: 1 pyruvate, 1 erythrose 4-phosphate (E4P) and 2 phosphoenolpyruvate (PEP).
14. **Arginine**: 5 ATP and 1 NADPH molecules are consumed. Amino acid precursors consumed are: 4 glutamate (glu), 1 glutamine (gln) and other metabolic precursors consumed are: 1 acetyl CoA (ac CoA), 2 bicarbonate and 1 pyruvate.
15. **Proline**: 1 ATP and 2 NADPH molecules are consumed. Amino acid precursor consumed is: 1 glutamate (glu).
16. **Lysine**: 2 NADPH molecules are consumed and 2 NADH molecules are produced. Amino acid precursors consumed are: 2 glutamate (glu) and other metabolic precursors consumed are: 1 acetyl CoA (ac CoA) and 1 2-oxoglutaric acid (2-OG).
17. **Histidine**: 1 ATP molecule is consumed and 2 NADH molecules are produced. Amino acid precursors consumed are: 1 glutamate (glu), 1 glutamine (gln) and other metabolic precursors consumed are: 1 5-phosphoribosyl pyrophosphate (PRPP).
18. **Leucine**: 1 NADPH molecule is consumed and 1 NADH molecule is produced. Amino acid precursor consumed is: 1 glutamate (glu) and other metabolic precursors consumed are: 1 acetyl CoA (ac CoA) and 2 pyruvate.
19. **Valine**: 1 NADPH molecule is consumed. Amino acid precursor consumed is: 1 glutamate (glu) and other metabolic precursors consumed are: 2 pyruvate.
20. **Isoleucine**: 3 ATP and 3 NADPH molecules are consumed. Amino acid precursors consumed are: 2 glutamate (glu) and other metabolic precursors consumed are: 2 pyruvate, 1 bicarbonate.

#### Biosynthetic cost calculation for the metabolic precursors

1. **Pyruvate**: 1 ATP and 1 NADH molecules are produced.
2. **3-phosphoglycerate**: 1 NADH molecule is produced.
3. **Phosphoenolpyruvate:** 1 NADH molecules are produced.
4. **Acetyl CoA**: 1 ATP and 2 NADH molecules are produced.
5. **5-phosphoribosyl pyrophosphate:** 2 ATP molecules are produced and 2 NADPH molecules are consumed.
6. **Erythrose 4-phosphate**: 2 ATP molecules are consumed.
7. **2-oxoglutaric acid**: 1 ATP and 4 NADH molecules are produced and 1 bicarbonate is consumed.
8. **5-methyl tetrahydrofolate**: 4 ATP and 3 NADPH molecules are consumed and 1 NADH molecule is produced. Amino acid precursors consumed are: 2 glutamate (glu), 1 glutamine (gln) and other metabolic precursors consumed are: 1 erythrose 4-phosphate (E4P), 2 phosphoenolpyruvate (PEP) and 1 3-phosphoglycerate (3-PG).
9. **Tetrahydrofolate**: 4 ATP and 2 NADPH molecules are consumed. Amino acid precursors consumed are: 1 glutamate (glu), 1 glutamine (gln) and other metabolic precursors consumed are: 1 erythrose 4-phosphate (E4P) and 2 phosphoenolpyruvate (PEP).

#### Final biosynthetic cost calculation for each amino acid including all its amino acid and metabolic precursors

For each amino acid, the total number of NADPH molecules consumed and the consolidated energetic cost in terms of high energy phosphate bonds (ATP and NADH) are listed in the Supplementary Table S1. For all the amino acid groups, the total number of NADPH molecules consumed and the consolidated energetic cost in terms of high energy phosphate bonds (ATP and NADH) are listed in the Supplementary Table S2.

